# Genomic evidence for ancient migration routes along South America’s Atlantic coast

**DOI:** 10.1101/2022.06.27.497820

**Authors:** Andre Luiz Campelo dos Santos, Amanda Owings, Henry Socrates Lavalle Sullasi, Omer Gokcumen, Michael DeGiorgio, John Lindo

**Author notes:** Correspondence (A.L.C.D.S.); (M.D.); (J.L.).

## Abstract

An increasing body of archaeological and genomic evidence has hinted to a complex settlement process of the Americas. This is especially true for South America, where unexpected ancestral signals have raised perplexing scenarios for the early migrations into different regions of the continent. Here we present ancient genomes from the archaeologically rich Northeast Brazil and compare them to ancient and present-day genomic data. We find a distinct relationship between ancient genomes from Northeast Brazil, Lagoa Santa, Uruguay and Panama, representing evidence for ancient migration routes along South America’s Atlantic coast. To further add to the existing complexity, we also detect greater Denisovan than Neanderthal ancestry in ancient Uruguay and Panama individuals. Moreover, we find a strong Australasian signal in an ancient genome from Panama. This work sheds light on the deep demographic history of eastern South America and presents a starting point for future fine-scale investigations on the regional level.

## Introduction

The Americas were the last continents populated by humans, with an increasing body of archaeological and genomic evidence indicating a complex settlement process starting from Beringia around the Last Glacial Maximum, ∼20,000 calendar years before present (BP) [1– 7]. Recent studies involving both ancient and present-day genomes have described how the ancestral Native Americans (NAs) further explored and settled northern North America and later diverged into two basal branches called Northern NA (NNA, or ANC-B) and Southern NA (SNA, or ANC-A) [3,4,7–11]. The SNA lineage, represented by the Clovis-associated Anzick-1 and the Spirit Cave individuals, is an ancestral component in present-day Central and South Americans, indicating that multiple groups related to this branch crossed Mesoamerica and entered South America. An additional nuanced ancestry to South America [1,7,8,10,11] may derive from an unsampled population (termed ‘Ypikuéra population’ or ‘Population Y’), which may have contributed to the early peopling of South America by introducing an Australasian shared ancestry that is observed in contemporary Indigenous Amazonian groups (e.g., Surui and Karitiana) [1,6,9,12,13]. To date, only one ancient individual (Sumidouro5, also one of the oldest representatives of the SNA lineage), unearthed in the Lagoa Santa archaeological area in Southeast Brazil, has been found to harbor the Australasian signal [7,9]. Vast portions of the southern continent, however, remain largely unexplored by archaeogenomic studies.

One such area is Northeast Brazil, along the Atlantic coast. Northeast Brazil houses some of the richest archaeological sites in South America [14–16] but has yielded only a single low-coverage ancient human genome to date (Enoque65, from Serra da Capivara archaeological area) [5] (supplemental information). In light of Brazil’s geographic extension, the archaeogenomic study of its Northeast Region may reveal important demographic aspects underlying many of the events that composed the settlement of South America, including putative migratory movements from North and Central America— through the Southern Cone along the Atlantic coast. The study of these still poorly characterized events from a genomic perspective, especially at the regional level, may lead to the disclosing of key chapters of the demographic history of the Americas [1,3,8,9].

Here, we report newly sequenced genomes from two ancient human individuals (Brazil-2 and Brazil-12) unearthed in two different archaeological sites in Northeast Brazil: Pedra do Tubarão and Alcobaça (Fig. 1A). Both archaeological sites are located in the state of Pernambuco and are associated with the Agreste rock art tradition, the second most representative rock art tradition in Northeast Brazil. It is believed that this tradition emerged at Serra da Capivara archaeological area, state of Piauí, approximately 5,000 years BP and later dispersed to other portions of Northeast Brazil. In Pernambuco, the oldest dates associated with this tradition go as far as 2,000 years BP. Brazilian archaeologists in Northeast Brazil have pointed to the challenges of affiliating rock art traditions and other material records at the archaeological sites in the area [17]. Thus, the chronological boundaries of putatively Agreste-affiliated archaeological cultures are still not precisely defined. There is also no record of post-European-contact Indigenous occupation of these sites, indicating a loss of cultural continuity in the area (supplemental information).

**Fig. 1.**
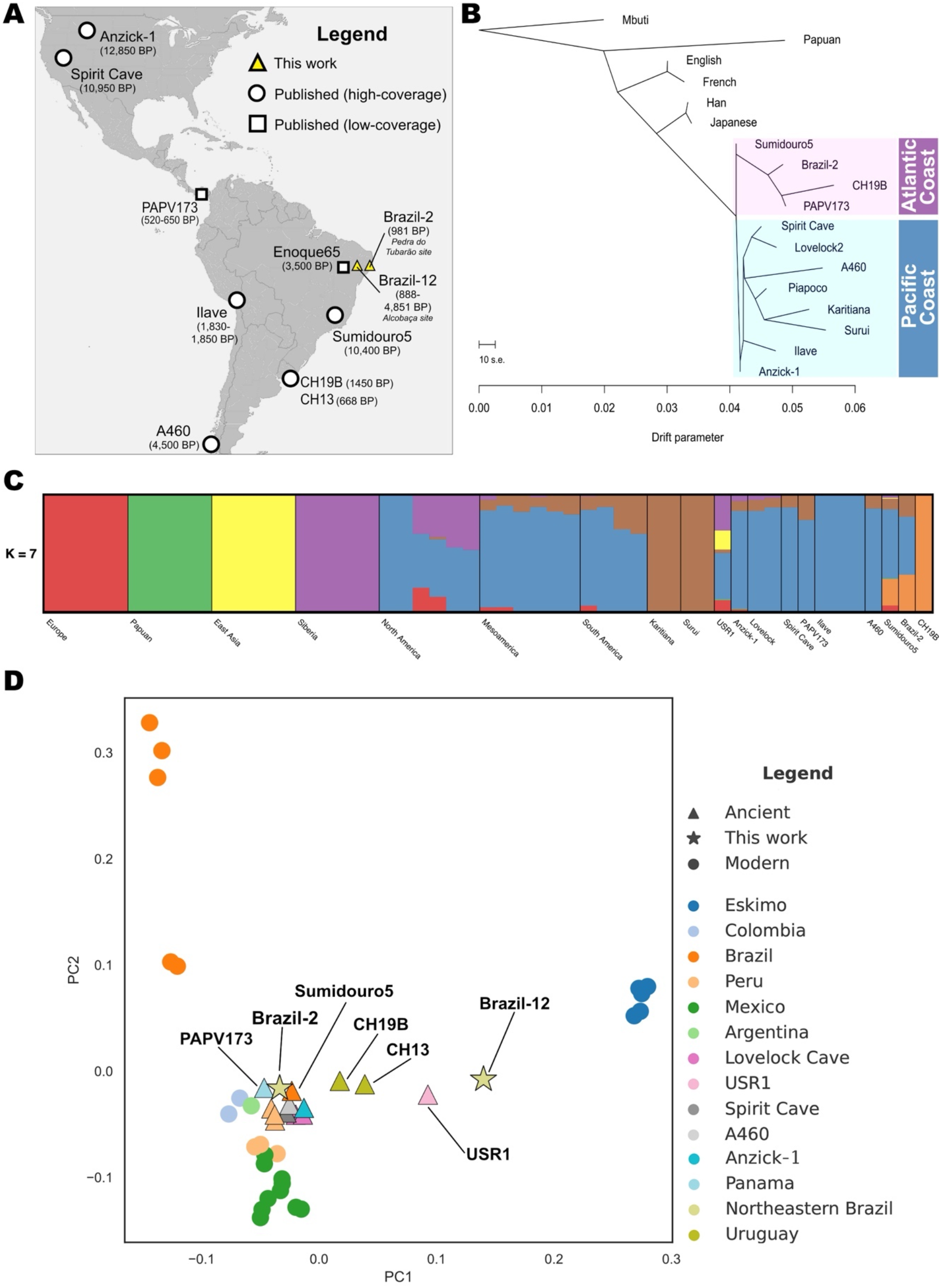
Ancient samples overview and broad genomic affinity. **(A)** Location of the newly sequenced ancient Brazilian samples and previously published ancient individuals. Ilave corresponds to individuals IL2, IL3 and IL7 from Peru. The chronological range associated with Brazil-12 represents contextual data (supplemental information). **(B)** TreeMix maximum likelihood tree showing the distinct Pacific and Atlantic clades highlighted. Only the ancient samples from Brazil and Uruguay with the highest sequencing depth were used. **(C)** ADMIXTURE analysis for *K*=7 clusters. Bars represent individuals, whereas each color represents a distinct ancestral component, with bar height representing the proportion of that component comprising a given individual. See fig. S1 for additional plots assuming different values for *K*. **(D)** Principal component analysis (PCA) on present-day individuals with ancient individuals projected onto the PCs 1 and 2. See fig. S2 for additional PCA plots.

The sequencing achieved a mean depth of approximately 10x for Brazil-2 and 8x for Brazil-12. Based on mitochondrial DNA, we estimated modern human contamination to be around 3.6% for Brazil-2 and 5% for Brazil-12. Brazil-2 has also been directly dated to approximately 981 years BP, with its molecular sex estimated to XY (supplemental information and Table 1). Along with the Northeast Brazil samples, we further investigated two recently sequenced ancient genomes from Uruguay (CH13 and CH19B) [18] (Fig. 1A). Our aim is to characterize, at the regional level, dispersal and admixture events involving the ancient individuals and populations of the Americas, particularly along South America’s Atlantic coast.

**Table 1.** Ancient individuals sequenced in this study and associated information. Radiocarbon dating of Brazil-2 was conducted at the University of Arizona AMS Laboratory. *Based on 24 traditional radiocarbon dates obtained from charcoals unearthed in the Alcobaça site. Therefore, the upper and lower boundaries presented here are, respectively, the mean values of the youngest and oldest ages obtained in the site (supplemental information).

| Sample | Site | <sup>14</sup>C Age (uncal. BP) | mtDNA Haplogroup | Contamination | Endogenous DNA | Mean Read Depth | Genetic Sex |
| --- | --- | --- | --- | --- | --- | --- | --- |
| **Brazil-2** | Pedra do Tubarão, Brazil | 981±23 | C1b | 3.6% | 11.65% | 10.67 | XY |
| **Brazil-12** | Alcobaça, Brazil | 888±25-4,851±30* | D1a2 | 5.0% | 3.41% | 8.32 | Not Assigned |

## Results

### A distinct genomic relationship between ancient Brazil, Panama and Uruguay

We investigated the ancient individuals’ broad genomic relationships to other populations using maximum likelihood trees [19], model-based clustering [20] and principal component analysis (PCA) [21] (Fig. 1, B to D) (Materials and Methods). We first evaluated the evolutionary relationship amongst ancient Native Americans (aside from the above-mentioned ancient genomes we also included the Andeans IL2, IL3 and IL7 from the Ilave region in Peru [22], the Chilean A460 [9] and PAPV173 from Panama [1]) and present-day worldwide populations via TreeMix [19]. The analysis separated the ancient Native Americans into two distinct clades (Fig. 1B). The first is composed of previously published samples unearthed near the Pacific coast of the Americas [9,22,23], in addition to the present-day Piapoco, Surui and Karitiana from the Amazonian rainforest [24]. The second clade comprises Sumidouro5 [9] and Brazil-2 from Brazil, CH19B [18] from Uruguay and PAPV173 from Panama [1]. While the first mentioned clade is representative of the ancient Americas’ Pacific coast, the second is composed of ancient individuals unearthed in archaeological sites closer to South America’s Atlantic coast, though we acknowledge that PAPV173 was found along Panama’s Pacific coast [1]. Still on the Atlantic clade, the resulting maximum likelihood tree indicates that Sumidouro5 is a possible ancestor of Brazil-2 (as it has no estimated genetic drift since diverging with Brazil-2), which in turn is associated with an ancestral branch of PAPV173 (and CH19B), a finding that suggests a south-to-north directionality. This result is consistent with the associated chronological data, i.e., Sumidouro5 is older than Brazil-2 and Brazil-2 is more ancient than PAPV173. On the other hand, the Pacific clade appears to summarize the body of knowledge [7] around the settlement of the Americas, in a north-to-south directionality.

We used ADMIXTURE to explore the genomic structure of the ancient Native Americans in the context of a worldwide reference panel [20]. When assuming *K* = 7 clusters, chosen based on the lowest cross-validation value, we found that Brazil-2, CH19B and Sumidouro5 share proportions of a distinct component, represented by the orange color. CH19B’s structure, specifically, is totally made up of the orange component. This ancestral component is restricted to ancient individuals unearthed closer to South America’s Atlantic coast. Interestingly, Sumidouro5 harbors a barely noticeable proportion of the green component, only found in present-day Papuans and in a few ancient samples from North America (USR1 and Anzick-1, also barely noticeable) (Fig. 1C and fig. S1).

Similarly, the PCA results show that Sumidouro5 falls between Brazil-2 and CH19B along PC1, with PAPV173 positioned very close to Brazil-2 (Fig. 1D and fig. S2). With the exception of the USR1 individual from Alaska, all other previously published ancient Native Americans are tightly clustered in proximity to Brazil-2, Sumidouro5 and PAPV173. This ancient cluster almost overlaps with present-day Peruvians and Argentinians, while present-day individuals from Mexico and Colombia are also in close proximity. At first glance, the clustering positions imply that Brazil-2 is more closely related to other present-day Native Americans than to the Karitiana and Surui from the Brazilian Amazonia. However, the observed distance between the ancient and present-day Brazilian samples is likely the result of strong genetic drift effects experienced by the Amazonian populations [6,12] (Fig. 1B) since they split with the ancient individuals. It is known that changes in allele frequencies due to drift in a given population might affect its position in PC-space [25] relative to other populations. The exception to this observation is when the referred population is not part of the PCs’ construction [25], which is not the case for Surui and Karitiana. In this context, outgroup *f*_3_ statistics (results provided in the *Surui and Karitiana harbor the highest affinity with ancient Americas* subsection further ahead) are a better suited [26] technique to assess the genomic affinity between the ancient and present-day Brazilian individuals. Brazil-12, on the other hand, falls closer to present-day Eskimo individuals than to any other present-day or ancient Native Americans, whereas CH13 (Uruguay) falls in the vicinity of USR1. It is important to note here that Brazil-12 and CH13 are shallow genomes compared to the other ancient individuals, which may explain the more distant clustering positions within the PCA plot.

### Deep archaic and Australasian ancestries in South America and Panama

We further explored the ancient individuals’ deep genomic ancestries using D-statistics [27] and identity by descent (IBD) analysis [28] (Fig. 2) (Material and Methods). We first evaluated the presence of the Australasian signal along South America’s Atlantic coast computing *D*-statistics of the form *D*(Yoruba,X;Mixe,TestPop), where X is a present-day non-African population and where TestPop is set as either Surui, Sumidouro5, PAPV173, Brazil-2, Brazil-12, CH19B, CH13 or Anzick-1. A similar *D*-statistic analysis was previously used to report this signal in the ancient Lagoa Santa genome (Sumidouro5) [9] and present-day Surui [6]. While we were able to replicate the Australasian signal in the Surui, we do not find the signal in Sumidouro5 as previously reported [9], possibly due to different reference panels being used (Fig. 2A). The signal is also not present in Brazil-2, Brazil-12, CH19B, CH13 and Anzick-1 (Fig. 2A and fig. S3). We do find, however, that Papuans, New Guineans and Indigenous Australians share significantly more alleles with PAPV173 than with the Mixe (Z > 3). Moreover, in the instances in which the Surui and PAPV173 were compared to the Mixe in relation to the Onge, a previously reported ‘attenuated signal’ [6] can be found (Z ≈ 2.7) (Fig. 2A). To corroborate these results, we tested some of the ancient samples using *D*-statistics of the form *D*(Yoruba,TestPop;X,B), in which B is set either as English, Han, Mixe, Papuan or Surui. Using this new form, we find that Sumidouro5 shares significantly more alleles with the Andamanese Onge (Z < −3) when B is the Papuans, which can represent the previously reported Australasian signal in this ancient sample [9] (table S1).

**Fig. 2.**
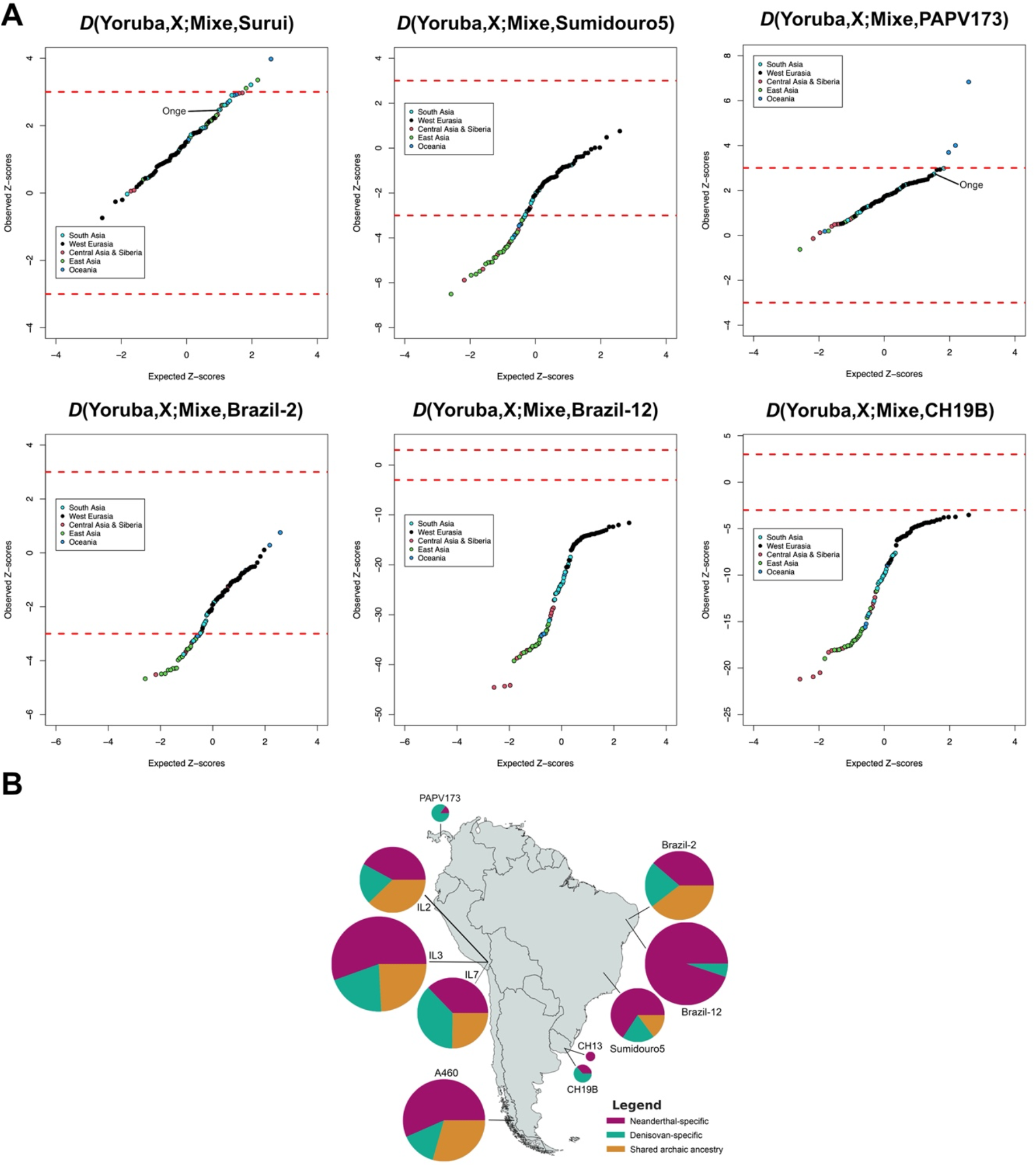
Deep ancestries of the ancient individuals of the Americas. **(A)** Quantile–quantile plots of the Z-scores for the *D*-statistic test for each ancient sample, a non-American, non-African population ‘X’ from Simons Genome Diversity Project (SGDP) [24], and the Mixe [24], compared to the expected ranked quantiles for the same number of normally distributed values. Each data point represents a distinct population from SGDP. Dashed red lines represent significance thresholds. **(B)** Archaic ancestry in ancient South America and Panama. Pie chart radius reflects the proportion of shared archaic loci in the individual.

To investigate even deeper genomic ancestry, we used IBDmix [28] to test all South American ancient individuals highlighted in this work (and the Panamanian PAPV173) for the presence of putative archaic (Altai Neanderthal [29] and Denisovan [30]) genomic contributions. We found that all samples share a very small genomic proportion with at least one of the archaic human species used as a reference (Fig. 2B). Interestingly, PAPV173 and CH19B harbor greater Denisovan-than Neanderthal-specific ancestry. When performing cluster analysis based only on the archaic proportions, these two samples cluster together (fig. S4), despite being situated more than 5,000 kilometers and almost 1,000 years apart, and is consistent with previous findings [18].

To corroborate the IBDmix results, we ran several *f*_4_-ratio tests to detect Denisovan-related ancestry in the high-coverage ancient samples: Brazil-2, IL2, IL3, IL7, Sumidouro5 and A460. We restricted our tests to the regions found to harbor archaic ancestry in the IBDmix analysis, i.e., the IBD tracks. All the non-African populations from SGDP public dataset were organized into super-/continental populations (Americas, Central Asia/Siberia, East Asia, South Asia and West Eurasia) and used here as baselines, i.e., we compare the proportion of Denisovan ancestry in the ancient individuals with the proportion harbored by the present-day superpopulations, one at a time (supplemental information). The resulting statistic (*α*) is defined as the ratio between two *f*_4_-statistics [31], and we use an already established *f*_4_-ratio form to test for the Denisovan-related ancestry [32,33] (supplemental information). Regardless of the baseline used, we find a positive correlation between the *f*_4_-ratio *α* and the proportion of Denisovan-related ancestry among the total archaic ancestry identified by IBDmix (fig. S5 and table S2). We recognize, however, that due to the small number of ancient samples tested here our results do not attain statistical significance.

### Surui and Karitiana harbor the highest affinity with ancient Americas

We used outgroup *f*_3_ statistics [31] to further highlight the shared genomic history between the ancient individuals and present-day populations (Fig. 3). Contrarily to what the PCA results implied (Fig. 1D and fig. S2), ranked outgroup *f*_3_ analyses demonstrate that Brazil2, Sumidouro5, PAPV173, CH19B, CH13 and Spirit Cave are more genomically related to Surui and Karitiana than to any other present-day population (Fig. 3 and fig. S6).

**Fig. 3.**
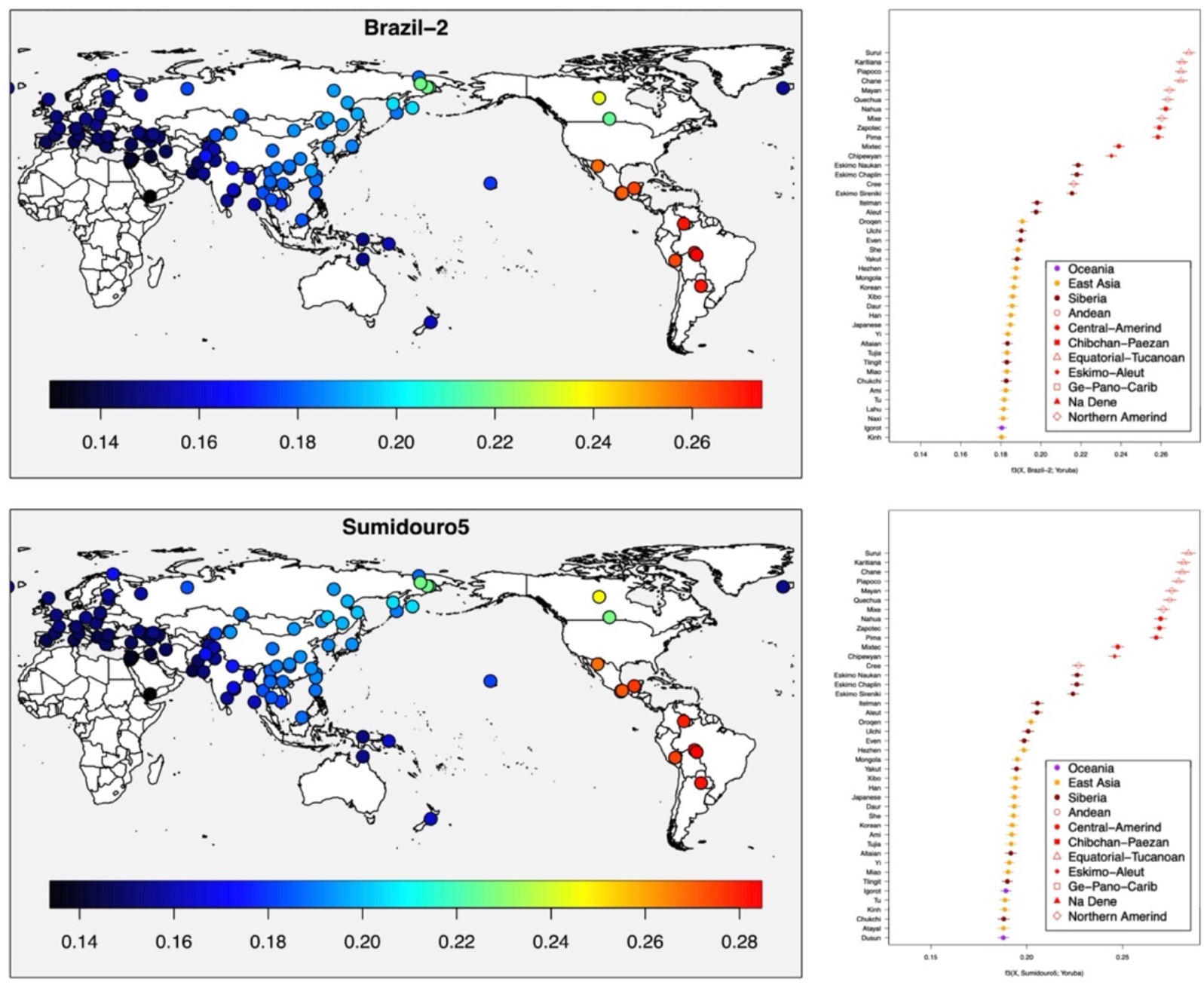
Top genomic affinities of ancient genomes from Brazil. Heatmaps and ranked results of outgroup *f*_3_-statistics analyses showing that Surui and Karitiana harbor the highest affinities with Brazil-2 and Sumidouro5, contrarily to what PCA implied (Fig. 1D).

### Dispersal in South America led to eastern two-way migration route

Lastly, we explored the demographic history of the ancient South American individuals by using demographic modeling information [31]. We used qpGraph [31] to build demographic models involving a reference panel of selected present-day worldwide populations and almost all the previously mentioned ancient individuals of the Americas (with the exceptions of Brazil-12 and CH13). The topology of the best-fit model, with three migration events, shows that population splits occurred after the first human groups reached South America’s western/Andean portion (as indicated by Quechua’s and Ilave’s position) (Fig. 4A). Brazil-2’s ancestry can be traced back both to a clade formed by Sumidouro5 and PAPV173, and to an ancestral branch of the present-day Piapoco, Surui and Karitiana. CH19B also received a big genomic contribution stemming from the clade formed by Sumidouro5 and PAPV173, while still inheriting genomic contribution from a possibly unsampled basal population, as previously reported [18] (Fig. 4a). Interestingly, the graph with two migration events shows that A460, Sumidouro5, Brazil-2 and PAPV173 form a clade by themselves, with CH19B receiving a large contribution from this clade, which is similar to the TreeMix result in Fig. 1B. This model suggests that the settlement of the Atlantic coast occurred only after the peopling of most of the Pacific coast (and the Andes). The Piapoco, Surui and Karitiana again form a distinct clade (Fig. 4B).

**Fig. 4.**
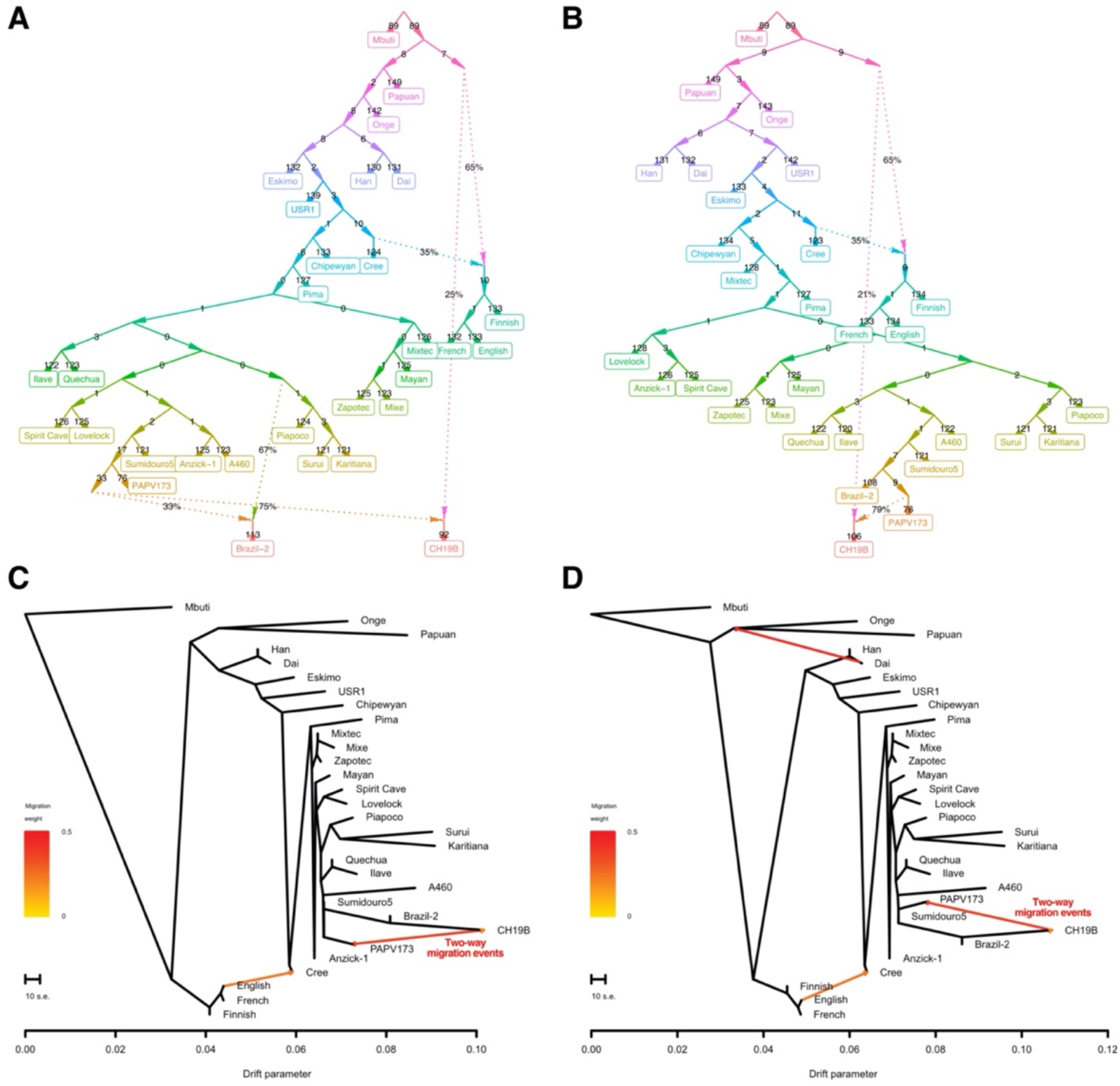
Demographic history of the ancient populations of South America. **(A)** and **(B)** Demographic models estimated by qpGraph with three and two migration events, respectively, showing populations splits in South America and the relationship between A460, Sumidouro5, Brazil-2, PAPV173 and CH19B. **(C)** and **(D)**, Demographic models estimated by TreeMix with three and four migration events, respectively, showing two-way migration events linking PAPV173 and CH19B.

Similarly, when maximum likelihood graphs involving the same samples/populations (and three and four migration events) are estimated using TreeMix, it is possible to observe Brazil-2, Sumidouro5 and CH19B forming a distinct clade, with PAPV173 and A460 as the nearest branches. These samples diverge only after the branching of an Andean clade formed by the Quechua and the Ilave ancient samples. Moreover, a two-way migration event linking CH19B and PAPV173 can be seen in both results (Fig. 4, C and D).

## Discussion

Consistent with previously-reported data [1], our results suggest that at least one population split likely occurred not long after the first SNA groups reached the southern portion of the Americas (Fig. 1B and Fig. 4, A and B). Based on the qpGraph results, we can hypothesize that this split took place around the Andes, later giving rise to ancient Southern Cone populations and the first groups that settled the Atlantic coast (Fig. 4, A and b, and Fig. 5). In light of Sumidouro5’s associated chronology—the oldest South American analyzed here—it is possible to affirm that the split occurred at least 10,000 years ago. Because Sumidouro5 is associated with the ancestors of both Brazil-2 and CH19B (Figs. 1B and 4), we can further conjecture that new migrations may have then emerged along the Atlantic coast, with Lagoa Santa as the putative geographical source of waves that headed in north-to-south and south-to-north directions—with the latter seemingly reaching Panama (Fig. 1B, Fig. 4, A and B, and Fig. 5). We conclude this hypothesis proposing that human movements closer to the Atlantic coast eventually linked Panama and Uruguay in a two-way migration route (Fig. 4, C and D, and Fig. 5). The migrations along the Atlantic coast apparently left no trace in the populations closer to the Pacific, as we could not find back-migration events in that direction (Fig. 4, C and D).

**Fig. 5.**
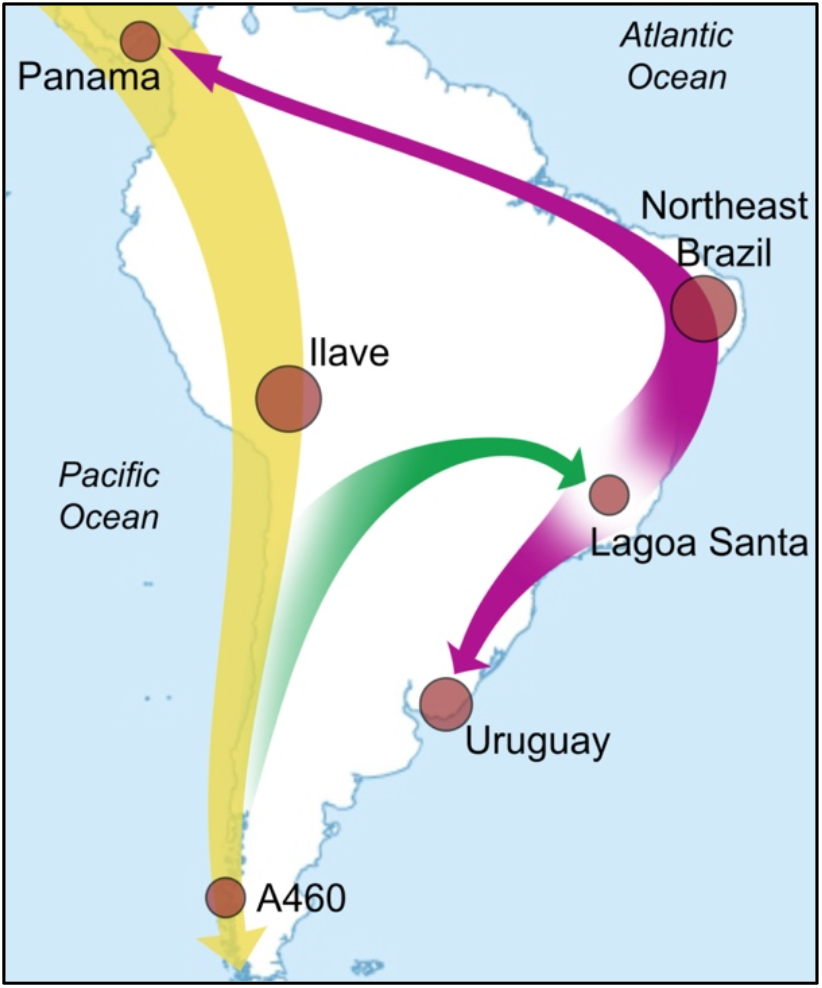
Hypothesis for ancient migrations in South America. The first SNA groups entered South America and spread through the Pacific coast settling the Andes (yellow arrow). At least one population split occurred soon after, branching the first groups that settled the Atlantic coast (green arrow) from the groups that gave rise to the ancient populations of Southern Cone. New migrations may have then emerged along the Atlantic coast, with a possible origin around Lagoa Santa, heading north toward Northeast Brazil and Panama, and south to Uruguay. Eventually, Uruguay and Panama were linked by a two-way migration route closer to the Atlantic coast (purple double-headed arrow).

Overall, our results show a strong genomic relationship among Brazil-2, CH19B, Sumidouro5 and PAPV173 (Fig. 1, B to D, and Fig. 4). Apart from the occurrence of mass burials in the sites that yielded these samples, there is no other evidence in the archaeological record that indicate shared cultural features between them. It is also important to note that Sumidouro5 is ∼9,000 years older than the other three mentioned individuals, enough time for expected and noticeable cultural divergence. Moreover, Brazil-2, CH19B and PAPV173, though more similar in age, were located thousands of kilometers apart from each other. Therefore, cultural differentiation is also expected among them [34]. On the other hand, our results further corroborate previously reported evolutionary relationships between PAPV173 and CH19B [18] by providing evidence of a distinct genomic affinity involving archaic human ancestry (Fig. 2B and fig. S4).

A strong signal of Australasian ancestry, previously reported only for the Lagoa Santa individual [9] and present-day Surui [6], was also observed for the previously published PAPV173, from Panama [1] (Fig. 2A). The Piapoco, Surui and Karitiana, however, harbor high affinities with Brazil-2 (Figs. 3 and 4A), and thus may have received contributions coming from Central America (in the form of the Australasian signal, in a north-to-south directionality) and South America’s Atlantic coast (in a south-to-north directionality).

Together, these results represent substantial genomic evidence for ancient migration events along South Americas’ Atlantic coast. Moreover, these events seem to have occurred as an outcome of the migratory waves that originated the first South American populations near the Pacific coast. With these findings we contribute to the unravelling of the deep demographic history of South America at the regional level.

## Author contributions

Conceptualization, A.L.C.D.S., H.S.L.S., O.G. and J.L.; Methodology, A.O. and J.L.; Investigation, A.L.C.D.S., J.L. and M.D.; Visualization, A.L.C.D.S., J.L. and M.D.; Supervision, J.L. and M.D.; Writing — Original Draft, A.L.C.D.S., J.L. and M.D.; Writing — Review & Editing, A.L.C.D.S., O.G., J.L. and M.D.

## Methods

### Experimental Design

Northeast Brazil harbors some of the richest and most diverse archaeological areas in the Americas [17], yet it remains largely unexplored by archaeogenomic studies. In light of Brazil’s geographic extension and position, genomic data of ancient individuals from the Northeast Region may reveal important aspects underlying many of the events that composed the settlement of South America along the Atlantic coast. We thus extracted and sequenced DNA from two archaeological individuals unearthed in Northeast Brazil to investigate ancient demographic and evolutionary aspects of the region. For this study, we employed established extraction and sequencing protocols (supplemental information), along with a variety of bioinformatics tools and statistical methods, such as: TreeMix [19], ADMIXTURE [20], PCA [21], *D*-statistics [6,27], IBDmix [28], *f*_4_-ratio [31], Outgroup *f*_3_ [31] and qpGraph [31].

### TreeMix Analysis

We started with the filtered dataset of called genotypes with transitions removed (supplemental information). TreeMix [19] was applied on the dataset to generate maximum likelihood trees and admixture graphs from allele frequency data. The Mbuti from the Simons dataset [24] was used to root the tree (with the ‘–root’ option). We accounted for linkage disequilibrium by grouping *M* adjacent sites (with the ‘–k’ option), and we chose *M* such that a dataset with *L* sites will have approximately *L*/*M* ≈ 20,000 independent sites. At the end of the analysis (i.e., number of migrations) we performed a global rearrangement (with the ‘–global’ option). We performed 20 iterations for each admixture scenario, choosing the best likelihood for each. We considered admixture scenarios with *m* = 0 and *m* = 3 migration events. A total of 726,182 overlapping sites were used.

### ADMIXTURE Analysis

We used the ADMIXTURE v.1.3 software to explore the genomic ancestral components in our dataset. The program computes a matrix of ancestral components proportions for each individual (Q) and provides a maximum likelihood estimate of allele frequencies for each ancestral component (P) [20]. Our dataset was investigated by specifying various numbers of hypothetical ancestral components (*K*). We ran ADMIXTURE assuming values from *K* = 2 to *K* = 8. The best run (i.e., the optimal value for *K*, with the most likely ancestral proportions) was selected based on the lowest tenfold cross-validation error after one hundred analysis iterations, by using the ‘--cv=10’ and ‘-C 100’ flags, respectively. After pruning the dataset for LD, 123,151 overlapping sites were utilized.

### Principal Component Analysis

PCA was performed using the ‘smartpca’ program from the EIGENSOFT v7.2.1 package [21]. The dataset used in this analysis integrated all 12 ancient genomes presented in Figure 1A plus USR1 [4] and Lovelock Cave [9] individuals, all with transitions removed, and 26 present-day individuals from the Simons Genome Diversity Project [24] (Chane from Argentina; Karitiana and Surui from Brazil; Piapoco from Colombia; Mayan; Mixe, Mixtec, Prima and Zapotec from Mexico; Quechua from Peru; and Eskimos Chaplin, Naukan and Sireniki from Russia). Principal components (PCs) were calculated using the present-day populations with the ‘poplistname’ and ‘autoshrink: YES’ options. Ancient data, characterized by a large portion of missing sites, were then projected onto the computed PCs with the ‘lsqproject: YES’ option. No outliers were excluded for this analysis, which was based on 2,727,376 loci presenting a genotyping rate of at least 90% across the whole dataset.

### D-statistics

The assessment of the Australasian signal in the ancient samples of the Americas and present-day Surui was performed using the POPSTATS Python program [6]. Two forms were used in this analysis. In the first analysis, we ran the analysis in the form of *D*(Yoruba,X;Mixe,TestPop), previously used to report the Australasian signal [6,9], in which ‘X’ was all non-African and non-American populations in the Simons Genome Diversity Project [24], while ‘TestPop’ was Brazil-2, Brazil-12, CH13 [18], CH19B [18], Surui [24], Sumidouro5 [9], Anzick-1 [23] or PAPV173 [1]. In the second analysis, we ran in the form of *D*(Yoruba,TestPop;X,B), in which ‘B’ was the English, Han, Mixe, Papuan and Surui populations from Simons Genome Diversity Project [24], ‘X’ was all the non-African populations from the same project [24] and ‘TestPop’ comprised the same samples used in the previous run, except for Anzick-1 [23] (supplemental information). The dataset had all transitions removed. No pruning for linkage disequilibrium was applied. The number of polymorphic sites used in this analysis depends on the coverage of the four populations that are being compared. The minimum number of sites analyzed was 143,014 sites in *D*(Yoruba,CH13;Chane,Surui) and the maximum was 3,831,654 sites in the *D*(Yoruba,Surui;Palestinian,Papuan).

### Identity by Descent Analysis

We assessed human archaic ancestry in the ancient samples of South America and Panama using the IBDmix software [28]. The program is able to identify introgressed human sequences using a pair of {‘archaic sample’}-{‘test population’} [28]. Since IBDmix needs at least 10 samples forming a single ‘test population’ to make robust inferences [28], we considered Brazil-2, Brazil-12, Sumidouro5 [9], CH13 [18], CH19B [18], A460 [9], IL2 [22], IL3 [22], IL7 [22] and PAPV173 [1] as a single population (‘Ancient NAs’). This ‘Ancient NAs’ dataset consisted of 5,010,609 polymorphic sites that were kept after transitions removal. No other filtering step was performed for this analysis. IBDmix was then run for each pair {Altai Neanderthal [29] or Denisova [30]}-{‘Ancient NAs’}. A summary of introgressed segments was generated and segments with a LOD score < 4 were filtered out with the ‘summary_lod: 4’ option. Introgressed segments with length < 50kb were also removed with the option ‘summary_length: 50000’. These are the same thresholds used in IBDmix’s original publication [28]. All the other parameters were run on default settings.

### *f*_4_-ratio Analysis

We performed *f*_4_-ratio tests to estimate Denisovan-related ancestry in the high-coverage ancient samples of South America using ADMIXTOOLS [31]. The genomes of Altai Neanderthal [29], Denisova [30] and all non-African (with the exception of Yoruba, which are used as an outgroup) populations in the Simons Genome Diversity Project [24] public dataset were compared using a previously established form [32,33] (supplemental information). The SGDP populations were then organized into superpopulations (Americas, Central Asia/Siberia, East Asia, South Asia and West Eurasia) and used as baselines, i.e., references for the amount of Denisovan-related ancestry [33]. The dataset had all transitions removed, and no pruning for linkage disequilibrium was applied. Only the genomic regions harboring archaic ancestry according to the IBDmix results were used. The number of polymorphic sites used in this analysis also depends on the coverage of the samples within the five populations that are being used in each test. The minimum number of sites analyzed was 1,630 when Sumidouro5 was tested, regardless of the baseline, whereas the maximum was 2,920 sites when IL3 was tested with the American superpopulation as baseline.

### Outgroup *f*_3_ Analysis

We extracted all non-African populations from the Simons Genome Diversity Project [24] as well as the sub-Saharan African Yoruba population to create a reference set of present-day human populations. For a given ancient sample (Sumidouro5, Brazil-2, CH19B, CH13, PAPV173, Anzick-1 or Spirit Cave), we merged variant calls from the ancient sample with the present-day human reference set. We then filtered for SNPs with exactly two distinct alleles observed in the merged set. To compute outgroup *f*_3_ statistics of the form *f*_3_(Present-day, Ancient; Yoruba) where the Yoruba population was considered the outgroup to the Present-day and Ancient human references, we applied the qp3Pop module of ADMIXTOOLS [31]. Because the Sumidouro5, Anzick-1 and Spirit Cave samples were not treated with uracil-DNA glycosylase, we also removed C/T and G/A SNPs to guard against a form of DNA damage. The number of polymorphic sites used in this analysis depended on the triple of populations being compared as well as whether C/T and G/A SNPs were removed from the samples. For the Sumidouro5, Anzick-1, and Spirit Cave samples in which we removed C/T and G/A SNPs, we respectively employed a minimum of 768,872, 763,566, and 778,916 sites and a maximum of 837,075, 829,379, and 848,782 sites. For the Brazil-2, CH19B, CH13, and PAPV173 samples, we respectively employed a minimum of 2,195,333, 737,689, 284,355, and 875,830 sites and a maximum of 2,363,461, 776,333, 297,652, and 932,752 sites.

### qpGraph

We extracted a subset of individuals from the Simons Genome Diversity Project [24] to create a small global reference panel to explore relationships between ancient (Brazil-2, Sumidouro5, PAPV173, CH19B, USR1, Anzick-1, Spirit Cave, Lovelock, IL2, IL3, IL7 and A460) and present-day samples using an admixture graph. The present-day populations we extracted were from Africa (Mbuti), Europe (Finnish, French and English), Oceania (Papuan), South Asia (Onge), East Asian (Han and Dai), Siberia (Eskimo Chaplin, Eskimo Naukan and Eskimo Sireniki that we jointly refer to as Eskimo in our analyses), North America (Cree, Chipewyan, Pima, Mixe, Mixtec and Zapotec), Central America (Mayan) and South America (Quechua, Piapoco, Karitiana and Surui). We merged this present-day reference panel with variant calls of the twelve ancients. We then filtered for SNPs with exactly two distinct alleles observed in the merged set, removed SNPs with any missing data, and removed C/T and G/A SNPs to guard against DNA damage, resulting in a dataset containing 110,505 SNPs. We applied the R package ADMIXTOOLS2 (https://uqrmaie1.github.io/admixtools/index.html, ADMIXTOOLS2 is currently under preparation) to perform qpGraph [31] estimation. Using this software, we precomputed *f*_2_ statistics between population pairs in a two megabase SNP block. Using a scenario with *M* ∈ {0,1,2,3,4} migration events, we initiated a graph search from a random initial graph with Mbuti set as the outgroup, and the algorithm for 1000 iterations. The graph search was rerun if the optimal graph with *M* migration events did not have a better score than those with fewer events. Comparing score distributions between 1000 bootstrap replicates of *f*_2_ blocks, we found that the best-fit model to have three migration events.

### Statistical Analysis

All statistical analyses employed in this work were previously implemented within the scope of the above-mentioned tools and methods.

### Data and materials availability

All data are available in the main text or the supplemental materials.

## References

1. Capodiferro MR et al. 2021 Archaeogenomic distinctiveness of the Isthmo-Colombian area. Cell 184, 1706–1723.e24. (doi:10.1016/j.cell.2021.02.040)

2. Flegontov P et al. 2019 Palaeo-Eskimo genetic ancestry and the peopling of Chukotka and North America. Nature 570, 236–240. (doi:10.1038/s41586-019-1251-y)

3. Lindo J et al. 2017 Ancient individuals from the North American Northwest Coast reveal 10,000 years of regional genetic continuity. Proc. Natl. Acad. Sci. U. S. A. 114, 4093–4098. (doi:10.1073/pnas.1620410114)

4. Moreno-Mayar JV et al. 2018 Terminal Pleistocene Alaskan genome reveals first founding population of Native Americans. Nature 553, 203–207. (doi:10.1038/nature25173)

5. Raghavan M et al. 2015 Genomic evidence for the Pleistocene and recent population history of Native Americans. Science 349, aab3884. (doi:10.1126/science.aab3884)

6. Skoglund P, Mallick S, Bortolini MC, Chennagiri N, Hünemeier T, Petzl-Erler ML, Salzano FM, Patterson N, Reich D. 2015 Genetic evidence for two founding populations of the Americas. Nature 525, 104–108. (doi:10.1038/nature14895)

7. Waters MR. 2019 Late Pleistocene exploration and settlement of the Americas by modern humans. Science 365. (doi:10.1126/science.aat5447)

8. Braje TJ, Dillehay TD, Erlandson JM, Klein RG, Rick TC. 2017 Finding the first Americans. Science 358, 592–594. (doi:10.1126/science.aao5473)

9. Moreno-Mayar JV et al. 2018 Early human dispersals within the Americas. Science 362. (doi:10.1126/science.aav2621)

10. Posth C et al. 2018 Reconstructing the Deep Population History of Central and South America. Cell 175, 1185–1197.e22. (doi:10.1016/j.cell.2018.10.027)

11. Scheib CL et al. 2018 Ancient human parallel lineages within North America contributed to a coastal expansion. Science 360, 1024–1027. (doi:10.1126/science.aar6851)

12. Silva MAC, Ferraz T, Bortolini MC, Comas D, Hünemeier T. 2021 Deep genetic affinity between coastal Pacific and Amazonian natives evidenced by Australasian ancestry. Proc. Natl. Acad. Sci. U. S. A. 118. (doi:10.1073/pnas.2025739118)

13. Skoglund P, Reich D. 2016 A genomic view of the peopling of the Americas. Curr. Opin. Genet. Dev. 41, 27–35. (doi:10.1016/j.gde.2016.06.016)

14. Boëda E et al. 2014 A new late Pleistocene archaeological sequence in South America: the Vale da Pedra Furada (Piauí, Brazil). Antiquity 88, 927–941. (doi:10.1017/S0003598X00050845)

15. Lahaye C et al. 2013 Human occupation in South America by 20,000 BC: the Toca da Tira Peia site, Piauí, Brazil. J. Archaeol. Sci. 40, 2840–2847. (doi:10.1016/j.jas.2013.02.019)

16. Santos GM, Bird MI, Parenti F, Fifield LK, Guidon N, Hausladen PA. 2003 A revised chronology of the lowest occupation layer of Pedra Furada Rock Shelter, Piauí, Brazil: the Pleistocene peopling of the Americas. Quat. Sci. Rev. 22, 2303–2310. (doi:10.1016/s0277-3791(03)00205-1)

17. Martin G. 1997 Pré-história do Nordeste do Brasil. Editora Universitária UFPE. See https://play.google.com/store/books/details?id=exmtOsvSKj4C.

18. Lindo J, De La Rosa R, dos Santos ALC, Sans M, DeGiorgio M, Figueiro G. 2022 The genomic prehistory of the indigenous peoples of Uruguay. PNAS Nexus (doi:10.1093/pnasnexus/pgac047)

19. Pickrell JK, Pritchard JK. 2012 Inference of population splits and mixtures from genome-wide allele frequency data. PLoS Genet. 8, e1002967. (doi:10.1371/journal.pgen.1002967)

20. Alexander DH, Novembre J, Lange K. 2009 Fast model-based estimation of ancestry in unrelated individuals. Genome Res. 19, 1655–1664. (doi:10.1101/gr.094052.109)

21. Patterson N, Price AL, Reich D. 2006 Population structure and eigenanalysis. PLoS Genet. 2, e190. (doi:10.1371/journal.pgen.0020190)

22. Lindo J et al. 2018 The genetic prehistory of the Andean highlands 7000 years BP though European contact. Sci Adv 4, eaau4921. (doi:10.1126/sciadv.aau4921)

23. Rasmussen M et al. 2014 The genome of a Late Pleistocene human from a Clovis burial site in western Montana. Nature 506, 225–229. (doi:10.1038/nature13025)

24. Mallick S et al. 2016 The Simons Genome Diversity Project: 300 genomes from 142 diverse populations. Nature 538, 201–206. (doi:10.1038/nature18964)

25. Peter BM. 2021 Modelling complex population structure using F-statistics and Principal Component Analysis. bioRxiv. (doi:10.1101/2021.07.13.452141)

26. Harris AM, DeGiorgio M. 2017 Admixture and ancestry inference from ancient and modern samples through measures of population genetic drift. Hum. Biol. 89, 21–46. (doi:10.13110/humanbiology.89.1.02)

27. Green RE et al. 2010 A draft sequence of the Neandertal genome. Science 328, 710–722. (doi:10.1126/science.1188021)

28. Chen L, Wolf AB, Fu W, Li L, Akey JM. 2020 Identifying and Interpreting Apparent Neanderthal Ancestry in African Individuals. Cell 180, 677–687.e16. (doi:10.1016/j.cell.2020.01.012)

29. Prüfer K et al. 2014 The complete genome sequence of a Neanderthal from the Altai Mountains. Nature 505, 43–49. (doi:10.1038/nature12886)

30. Meyer M et al. 2012 A high-coverage genome sequence from an archaic Denisovan individual. Science 338, 222–226. (doi:10.1126/science.1224344)

31. Patterson N, Moorjani P, Luo Y, Mallick S, Rohland N, Zhan Y, Genschoreck T, Webster T, Reich D. 2012 Ancient admixture in human history. Genetics 192, 1065–1093. (doi:10.1534/genetics.112.145037)

32. Qin P, Stoneking M. 2015 Denisovan ancestry in east Eurasian and Native American populations. Mol. Biol. Evol. 32, 2665–2674. (doi:10.1093/molbev/msv141)

33. Carlhoff S et al. 2021 Genome of a middle Holocene hunter-gatherer from Wallacea. Nature 596, 543–547. (doi:10.1038/s41586-021-03823-6)

34. Eerkens JW, Lipo CP. 2007 Cultural transmission theory and the archaeological record: Providing context to understanding variation and temporal changes in material culture. J. Archaeol. Res. 15, 239–274. (doi:10.1007/s10814-007-9013-z)

