## supplemental information for "Genomic evidence for ancient migration routes along South America’s Atlantic coast"

**This file includes:**

Supplementary Text
Figs. S1 to S7
Tables S1 to S3

**Other Supplementary Materials for this manuscript include the following:**

Data S1

**Supplementary Text**

Archaeological context

Northeastern Brazil has almost 50% of its geographical space occupied by hot and semi-arid lands (with summer rains), since it is located far from the two largest drainage basins in South America (Amazon and Rio de la Plata) [1,2]. The oldest prehistoric settlement pathways in the region are still unknown, as the current state of knowledge does not allow for affirmations with solid scientific bases. There is no evidence of prehistoric occupations along the Atlantic coast in the Pleistocene, a fact that can be explained by the increasing sea levels since the ending of the Last Glacial Maximum (LGM), which may have destroyed possible signs of ancient human occupations in Northeastern Brazil. The known archaeological coastal occupations belong, for the most part, to ceramist groups. The oldest archeological records are found mainly in the karst formations, which indicates that the human groups that inhabited the region sought out deep limestone shelters to protect themselves [2]. Within the expected cultural variability, there is, however, an indisputable ancestral homogeneity in the different Brazilian human groups, which, along with all other South American human groups, are associated with a single genomic lineage [2–5].

The largest and richest archeological area in Northeast Brazil, and probably in the Americas, is the Serra da Capivara National Park, located in the state of Piauí [2,6–8]. Brazilian archaeologists captained by Niède Guidon, Anne-Marie Pessis and Gabriela Martin have documented the existence of more than 1,000 archaeological sites in the area in the last five decades, including the largest concentration of rock paintings in the American continents [2,8]. Regarding the prehistoric human skeletal remains in the area, prior to this work, the sample named Enoque65, unearthed in the 'Toca do Enoque' archaeological site, is the only human remain from Serra da Capivara and Northeastern Brazil to be ever sequenced and analyzed in an archaeogenomics study to date [9]. In cultural terms, the Serra da Capivara archaeological area is the epicenter of rock art traditions that later dispersed to other areas within Northeast Brazil [10]. The second most representative rock paintings in the region belong to the 'Agreste' tradition, which is believed to be have emerged in Serra da Capivara approximately 5,000 years BP. Specimens of this traditions were documented in the two sites that yielded samples to this work: Pedra do Tubarão and Alcobaça, located in the state of Pernambuco [2,11]. There, the Agreste tradition is associated with dates around 2,000 years BP. However, according to Brazilian archaeologists, affiliating rock art traditions and other material remains at the archaeological sites in the area is a challenging task. Therefore, the chronological boundaries of archaeological cultures putatively affiliated with the Agreste tradition remains to be precisely defined. There is also no evidence of Indigenous occupation of these sites after the European contact, indicating a loss of cultural continuity in the area [2].

Pedra do Tubarão

Pedra do Tubarão (Shark Rock) archaeological site, is a rock shelter located in the municipality of Venturosa (latitude: 8º32'21'' S; longitude: 36º48'08'' W), state of Pernambuco, Northeast Brazil, and was excavated in 1989 by a team of archaeologists from the Federal University of Pernambuco (UFPE). The rock shelter is characterized by Agreste-affiliated panels covering big portions of its area. Two distinct prehistoric periods of human occupations have been identified in the site: a more ancient occupation characterized by the use of lithic implements; and another one, more recent, defined by the use of pottery. However, no chronological boundaries have been estimated for these prehistoric occupations [2]. An indigenous burial ground, named Cemitério do Caboclo (Caboclo Graveyard), was found at approximately 200 meters from the site and was also excavated. The skeletal remains of at least 24 ancient human individuals were unearthed there and are currently stored in the Biological and Forensic Archaeology Laboratory (LABIFOR) at UFPE. Purposely broken and burned human bones and mass burials are products of the funeral ritual characteristic of this site [2]. Prior to this work, three bone samples have been directly dated to ~1,000 BP [2,12]. A tooth specimen collected in this site was sequenced and presented here as Brazil-2. This specimen was found unarticulated and thus cannot be associated with a specific individual buried at the site (consequently, it is also not possible to estimate the age at death for this individual). A sample of this specimen have been dated to 981 calendar years BP at the University of Arizona AMS Laboratory. However, the lack of contextual chronological data for the site makes it not possible to affiliate this tooth to a specific occupation.

<u>Alcobaça</u>

The Alcobaça archaeological site is located in the municipality of Buíque (latitude: 8°32'24'' S; longitude: 37°11'39'' W), state of Pernambuco, Northeast Brazil. A 10-meters-high rock shelter characterized by rich Agreste-affiliated panels covering large portions of its area, the site was also excavated by a team of archaeologists from UFPE between 1996 and 1998, yielding five human burials. These remains are also currently stored in the LABIFOR at UFPE. Similar to the Pedra do Tubarão archaeological site, some of the bony remains were purposely burned and presented red painting, the predominant color in the rock art panels [2]. The site's soil presents a highly thick layer of ashes, which allowed for 24 traditional radiocarbon dates from charcoal unearthed there, providing a chronological sequence for intense human occupation that ranged from 4,851±30 BP to 888±25 BP [2,12]. Iron oxide used to create the paintings and pottery fragments were also found in the site [2]. A tooth specimen associated with 'burial 2' was sequenced and presented in this work as Brazil-12. Burial 2 was also composed of burned, unarticulated skeletal remains, and no analysis regarding ages at death were performed on the bones. The chronological data obtained for this site seems to suggest the final periods of the Agreste tradition and associated archaeological records/cultures [2].

<u>Ancient DNA and Sequencing</u>

Samples were extracted for each ancient tooth, and sequencing libraries were constructed at the Lindo Ancient DNA laboratory at Emory University using the Dabney protocol [13]. Libraries were prepared with the NEB Ultra II DNA Library Prep for Illumina, with modifications for ancient DNA, which included quartering the reagents, the utilization of 1:20 adaptor dilution, and 1.5 ul of premixed NEB unique dual indexes. Samples were preliminarily screened for endogenous DNA on the Illumina iSeq 100 at the Lindo Lab, with libraries that were not treated with the USER enzymes. Samples that were selected for deep sequencing on the NovaSeq 6000 at Dante Labs (L'Aquila, Italy), included libraries treated with the USER enzyme to help compensate for DNA damage.

The ancient raw sequences were trimmed for Illumina adapters using AdapterRemoval2 [14] and aligned to the hg19 human reference sequence using the BWA mem algorithm [15], which has been shown to increase accuracy with ancient DNA mapping over the *aln* algorithm [16]. The alignments from the shotgun sequences that were not treated with the USER enzyme were used to validate their ancient authenticity with MapDamage2 [17]. The two ancient individuals from Brazil showed deamination patterns consistent with ancient DNA (fig. S7). The mean read depths of the assemblies were calculated using the program Mosdepth [18]. Whereas the molecular/genetic sex of Brazil-2 was estimated using the ry_compute Python program [19] (Table 1)—Brazil-12's genetic sex could not be estimated due to overall lower coverage.

Genotypes were called from both ancient individuals using the Ancient DNA Caller ARIADNA, which uses a machine learning method to overcome issues with DNA damage and contamination [20]. The resulting VCF file was further filtered to remove genotype calls with allele counts below 3 and for sites that violated a one-tailed test for Hardy–Weinberg Equilibrium at a p-value of $10^{-4}$ with VCFtools [21]. The dataset was then merged with modern and ancient samples from the Americas with bcftools [21,22]. To guard against potential bias caused by post-mortem DNA damage patterns, characteristic of ancient individuals, we also excluded the C→T and G→A transition polymorphic sites from the merged VCF file. Mitochondrial and allosomic data were also filtered out.

All of the ancient reference samples were also called in the same way using the ARIADNA Ancient DNA caller, to prevent batch affect. This includes the reference samples Spirit Cave, Lovelock2, Lovelock3, Anzick-1, Llave, Chile_A460, Sumidouro5, and CH19B. PAPV173, a low coverage sample, was also called with ARIADNA, since the method has been shown to effectively remove biased heterozygous sites resulting from noisy low-coverage data.

Contamination estimates

Contamination was assessed using the mitochondrial genomes of all three ancient individuals unearthed in Brazil. We ran the software Schmutzi [23] for contamination estimation in each sample (Table 1). This software not only estimates present-day human contamination in the sample but also reconstructs the assessed endogenous mitochondrial genome by considering both deamination patterns and fragment length distributions [24].

To corroborate the results of Schmutzi [23], we also ran the program Haplocheck 1.3.3 [25], which only detected contamination in Brazil-2, however, in a similar proportion as estimated by Schmutzi (table S3).

Corroborating the initial D-statistics results

The first round of *D*-statistics [26] runs, of the form *D*(Yoruba,X;Mixe,TestPop) [27], revealed that Papuans, New Guineans and indigenous Australians share significantly more derived alleles with PAPV173 than with the Mixe ($Z > 3$), when X is Papuan, Bougainville or Australians, respectively, and TestPop is PAPV173. While we find this previously unreported signal in PAPV173, we were unable to replicate a previously reported Australasian signal in Sumidouro5 [4].

To corroborate these results, we tested some of the ancient samples using a different form, *D*(Yoruba,TestPop;X,B), in which B could be English, Han, Mixe, Papuans or Surui. Whereas TestPop was Brazil-2, Brazil-12, PAPV173, Sumidouro5, CH19B, CH13 and Surui. In this new form, we find that Sumidouro5 share significantly more alleles with the Andamanese Onge ($Z < -3$) when B is Papuan (table S1). This result seems to replicate the previously reported signal in Sumidouro5 [4].

Corroborating the IBDmix results

To corroborate our IBDmix [28] findings we used ADMIXTOOLS [29] to run several f4-ratio statistics tests for the ancient samples and different baselines. The Denisova-related [30] ancestry was calculated directly, using a similar form as in Qin and Stoneking (2015) [31] and Carlhoff *et al*. (2021) [32] (Extended Data Fig. 5a):

$$\alpha = \frac{f_4(\text{Yoruba, Altai Neanderthal; Ancient, Baseline})}{f_4(\text{Yoruba, Altai Neanderthal; Denisova, Baseline})}$$

in which $\alpha$ is the Denisova-related signal, and 'Ancient' were all the high-coverage samples from South America analyzed with IBDmix [28]: Brazil-2, Sumidouro5 [4], IL2, IL3, IL7 [24] and A460 [4]. The baselines used were the superpopulations from Simons Genome Diversity Project [33]: Americas (AMR), Central Asia/Siberia (CAS), East Asia (EAS), South Asia (SAS) and West Eurasia (WEU)—with the exception of the Yoruba, which was used as an outgroup instead, alongside the Altai Neanderthal genome [34].

The results show that, regardless of the baseline superpopulation used in the tests, we find a positive correlation between estimates of $\alpha$ and the proportion of Denisovan-related ancestry among the total archaic ancestry identified with IBDmix [28] (fig. S5 and table S2).

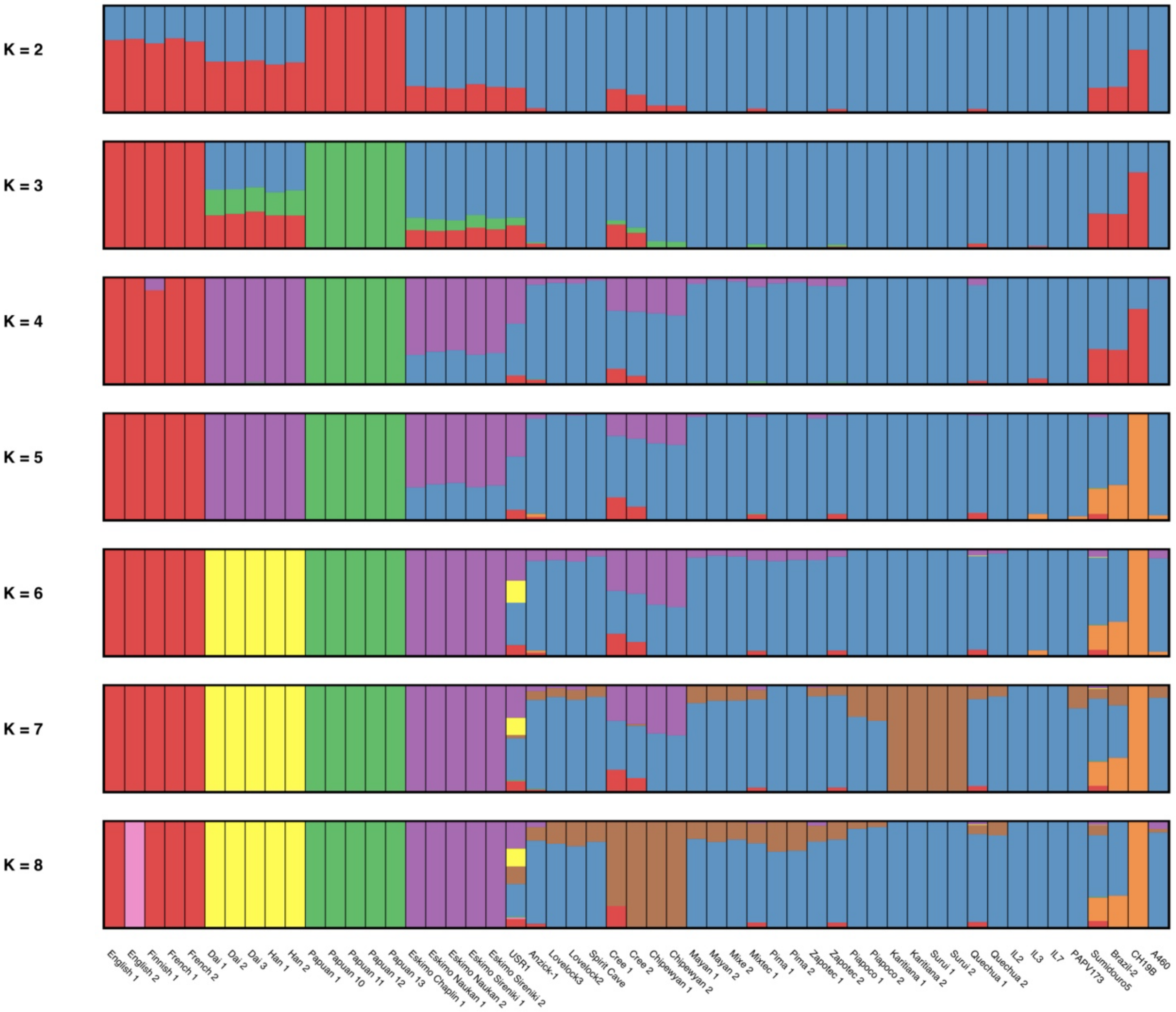


**Fig. S1. ADMIXTURE results assuming values for *K* equal to 2 through 8.** Bars represent individuals, whereas each color represents a distinct ancestral component, with bar height representing the proportion of that component comprising a given individual.

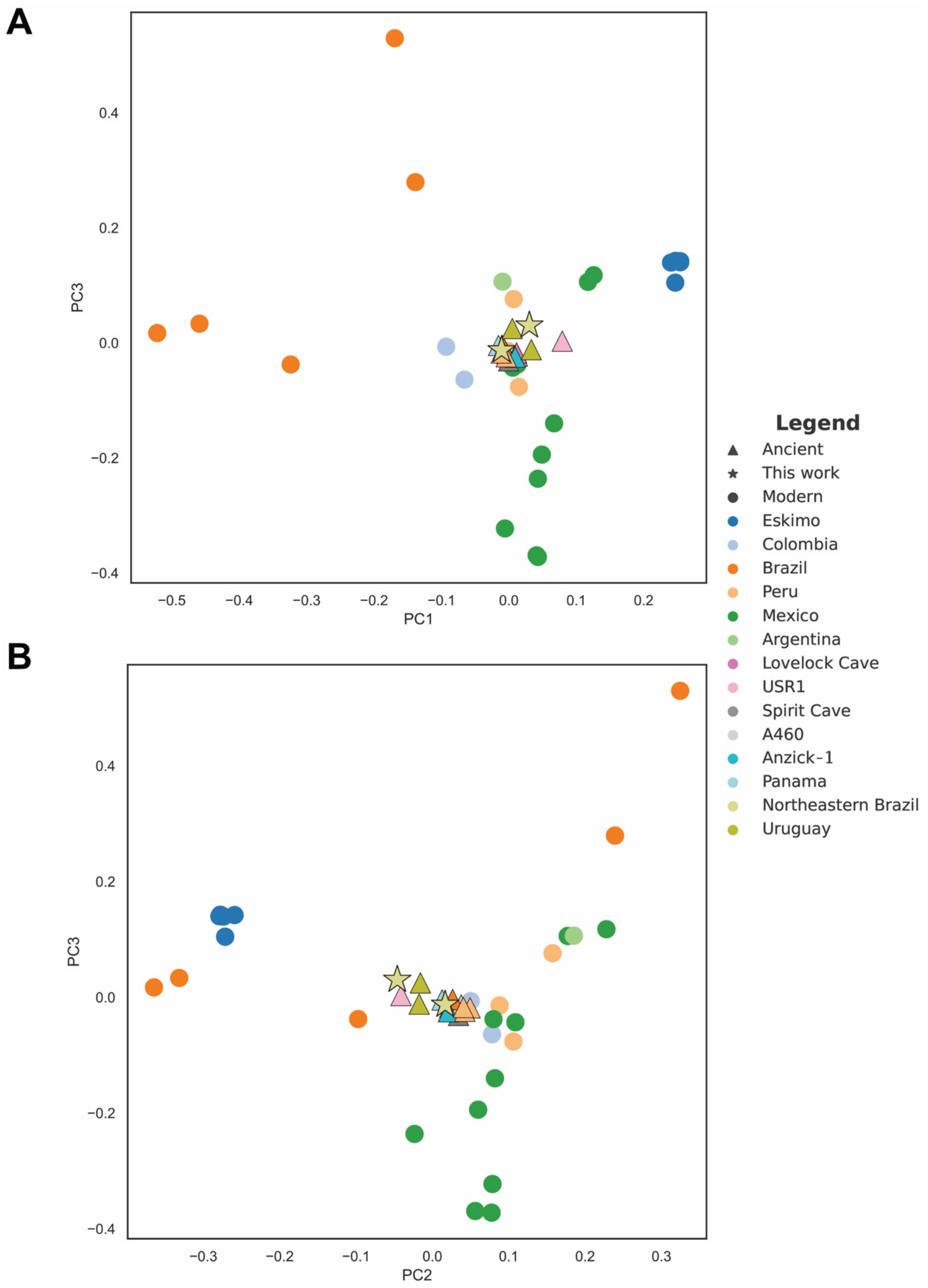


**Fig. S2. Alternative principal component analysis (PCA) plots with samples unlabeled. (A)** PC1 versus PC3. **(B)** PC2 versus PC3. PCA was applied to present-day individuals with ancient individuals projected onto the identified components.

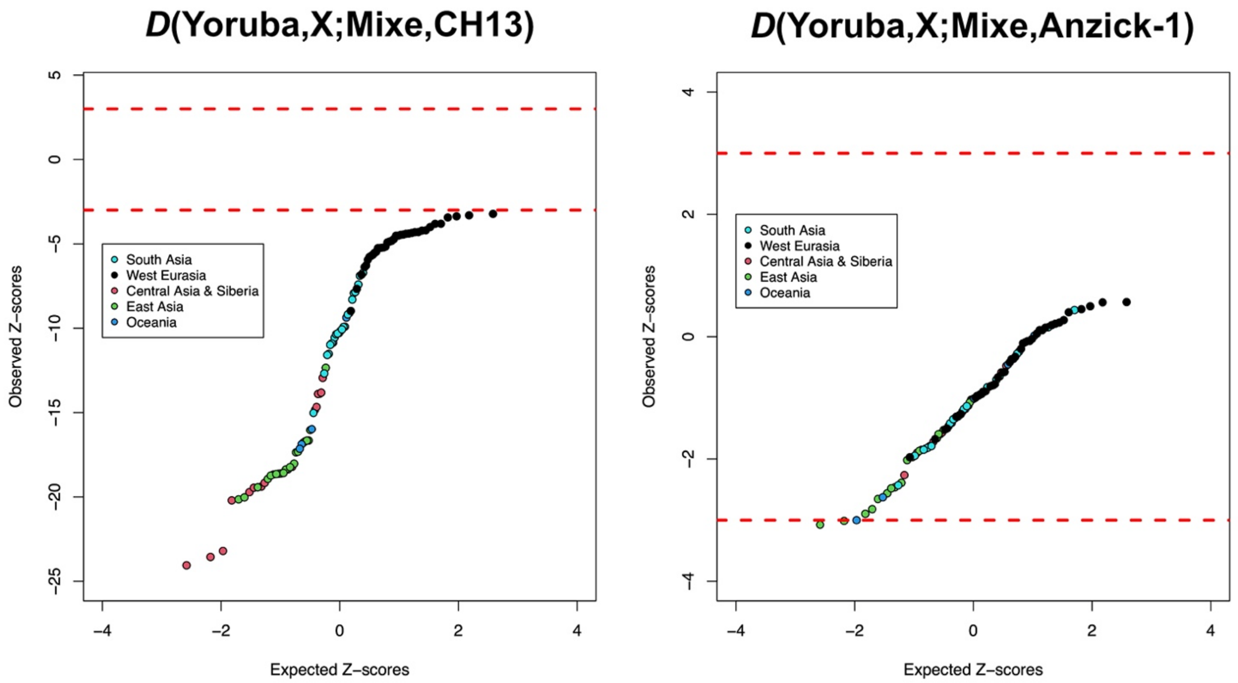


**Fig. S3. Quantile–quantile plots of the Z-scores for the *D*-statistics for CH13 and Anzick-1, compared to the expected ranked quantiles for the same number of normally distributed values.** Each data point represents a distinct population. Dashed red lines represent significance thresholds.

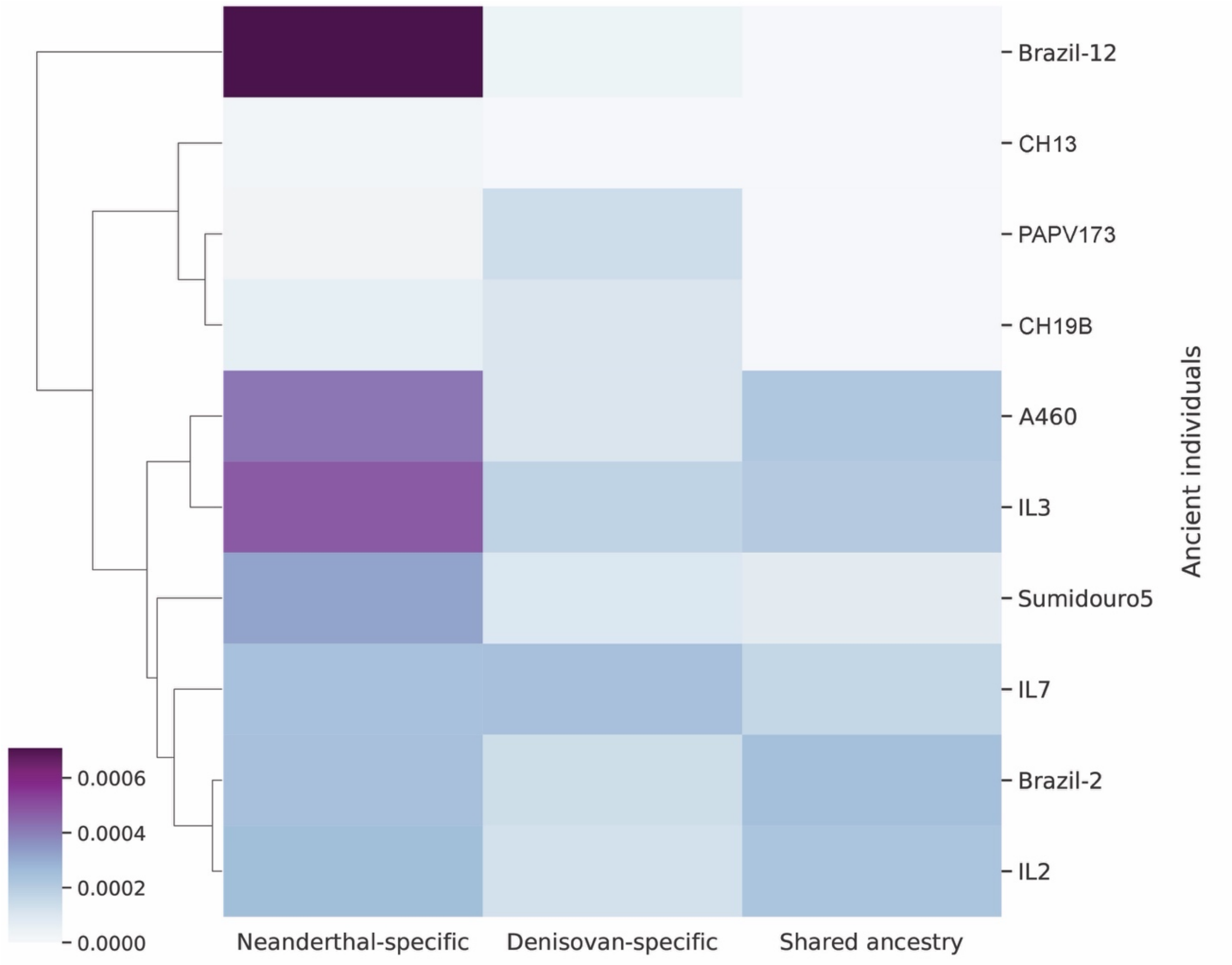


**Fig. S4. Proportions of archaic hominin ancestry in ancient South America and Panama.** This hierarchically clustered heatmap shows that the Panamanian PAPV173 and the Uruguayan CH19B cluster together mainly because they harbor higher proportions of Denisovan-specific than Neanderthal-specific ancestry, despite being situated more than 5,000 kilometers and almost 1,000 years apart. The colorbar represents the proportion of archaic hominin ancestry.

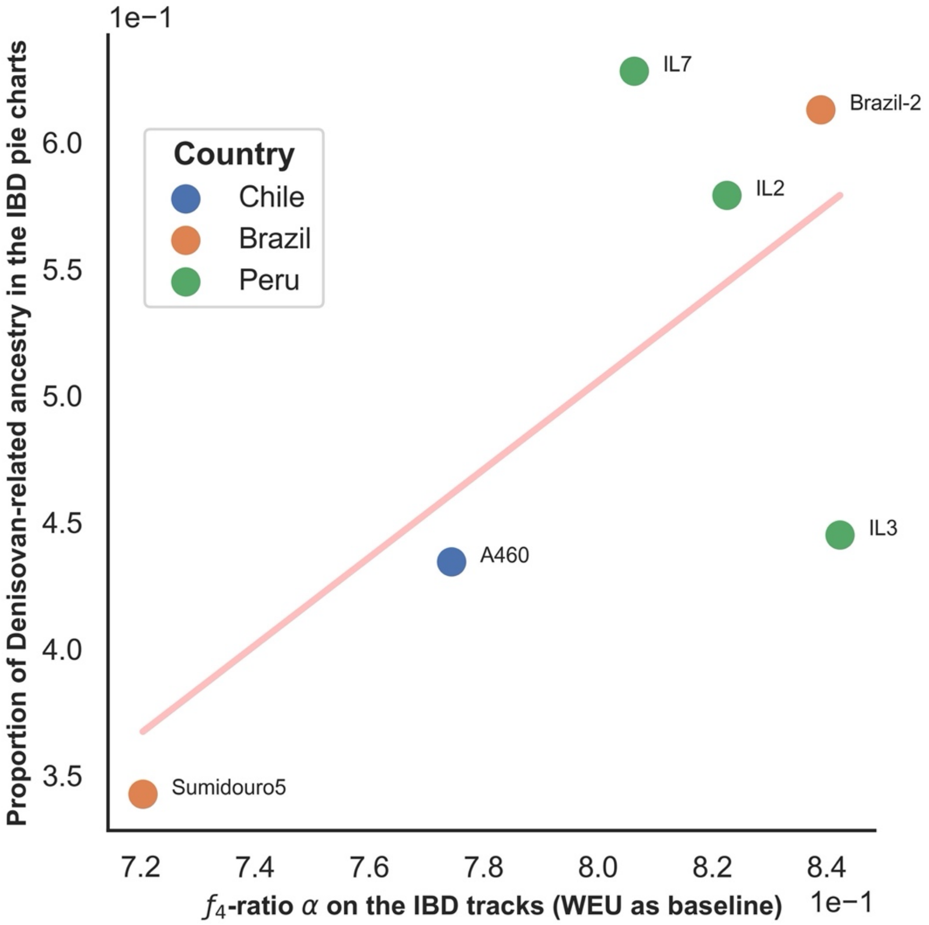


**Fig. S5. Correlation between the $f_4$-ratio and IBDmix results for the Denisovan-related ancestry.** Positive correlation between $f_4$-ratio $\boldsymbol{\alpha}$ (West Eurasians as baseline) and IBDmix results. See supplementary text for more details on the $f_4$-ratio tests and correlation tests using different baselines.

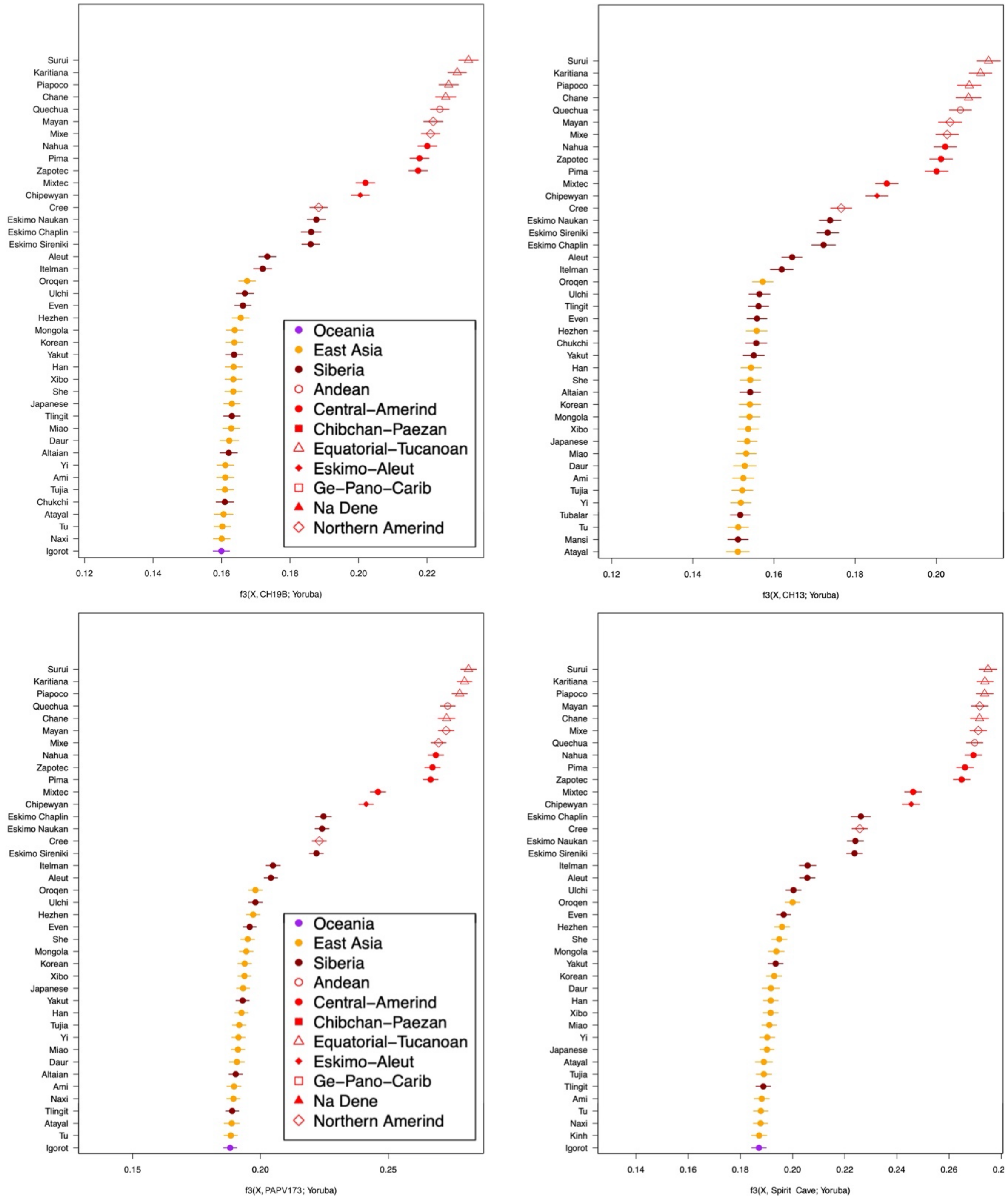


**Fig. S6. Ranked Outgroup *f*$_3$-statistics' results for CH19B, CH13, Spirit Cave and PAPV173 (clockwise).** Surui and Karitiana harbor the highest affinities with these ancient samples.

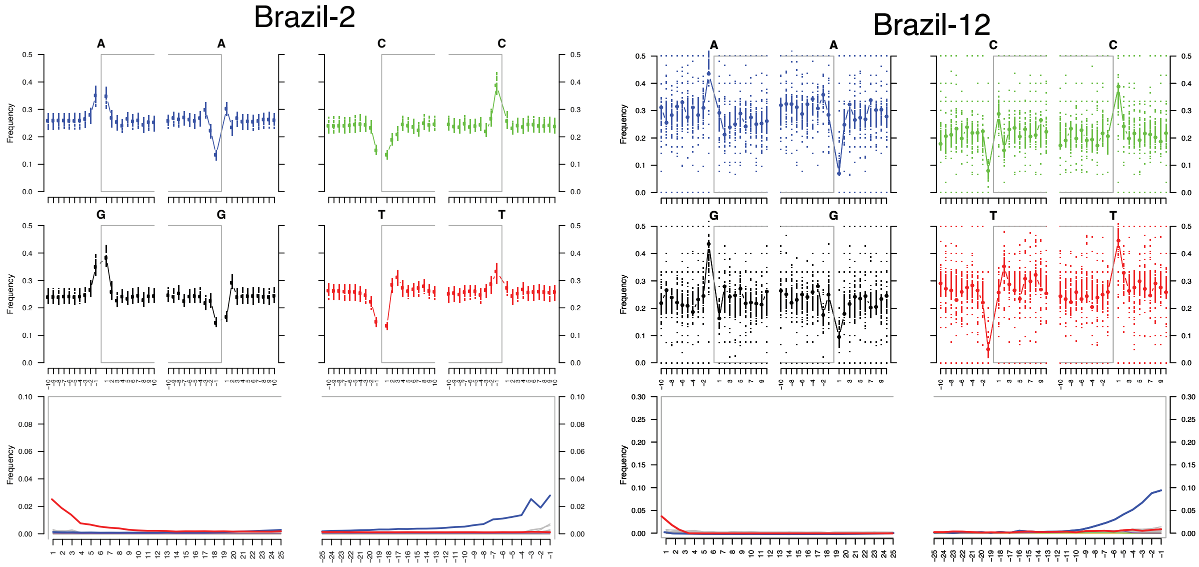


**Fig. S7. MapDamage graphs for the two ancient individuals from Brazil sequenced in this study.**

**Table S1. *Z*-scores for *D*-statistics tests of the form *D*(Yoruba,TestPop;Onge,B).** Highlighted Z-scores show that PAPV173 and Sumidouro5 share significantly more alleles with the Andamanese Onge ($Z < -3$), when B is English and Papuan, respectively. Specifically, PAPV173 and Surui share a significant Onge signal when compared to the English.

| Outgroup | TestPop | X | B | *Z*-score |
|---|---|---|---|---|
| Yoruba | Brazil-2 | Onge | English | -2.6469731 |
| Yoruba | Brazil-2 | Onge | Han | 17.0306987 |
| Yoruba | Brazil-2 | Onge | Mixe | 48.8807101 |
| Yoruba | Brazil-2 | Onge | Papuan | -1.9257698 |
| Yoruba | Brazil-2 | Onge | Surui | 53.6195854 |
| Yoruba | Brazil-12 | Onge | English | 9.2697455 |
| Yoruba | Brazil-12 | Onge | Han | 6.8578083 |
| Yoruba | Brazil-12 | Onge | Mixe | 18.523200 |
| Yoruba | Brazil-12 | Onge | Papuan | -1.3633319 |
| Yoruba | Brazil-12 | Onge | Surui | 20.8161594 |
| Yoruba | PAPV173 | Onge | English | -3.8574129 |
| Yoruba | PAPV173 | Onge | Han | 17.5773349 |
| Yoruba | PAPV173 | Onge | Mixe | 52.1726797 |
| Yoruba | PAPV173 | Onge | Papuan | -1.9429649 |
| Yoruba | PAPV173 | Onge | Surui | 53.2342001 |
| Yoruba | Sumidouro5 | Onge | English | -2.7235643 |
| Yoruba | Sumidouro5 | Onge | Han | 14.6953653 |
| Yoruba | Sumidouro5 | Onge | Mixe | 41.6379945 |
| Yoruba | Sumidouro5 | Onge | Papuan | **-3.9313316** |
| Yoruba | Sumidouro5 | Onge | Surui | 42.8940762 |
| Yoruba | CH19B | Onge | English | 1.08750147 |
| Yoruba | CH19B | Onge | Han | 14.5992294 |
| Yoruba | CH19B | Onge | Mixe | 43.6985309 |
| Yoruba | CH19B | Onge | Papuan | -1.8303643 |
| Yoruba | CH19B | Onge | Surui | 47.9261463 |
| Yoruba | CH13 | Onge | English | 1.72084862 |
| Yoruba | CH13 | Onge | Han | 12.0119818 |
| Yoruba | CH13 | Onge | Mixe | 36.3286196 |
| Yoruba | CH13 | Onge | Papuan | -2.1094606 |
| Yoruba | CH13 | Onge | Surui | 38.5810098 |
| Yoruba | Surui | Onge | English | -4.99601930 |
| Yoruba | Surui | Onge | Han | 18.7136901 |
| Yoruba | Surui | Onge | Mixe | 53.0279858 |
| Yoruba | Surui | Onge | Papuan | -1.0693530 |

**Table S2. Positive correlation between $f_4$-ratio $\alpha$ and IBDmix results.**

| **Superpopulation** | **Pearson's *r*** |
|---|---|
| AMR | 0.6885467 |
| CAS | 0.6843622 |
| EAS | 0.68487689 |
| SAS | 0.69634196 |
| WEU | 0.69777823 |

**Table S3. Contamination estimates for the newly sequenced samples from Northeast Brazil.** Both Schmutzi and Haplocheck estimated similar contamination values for Brazil-2. Haplocheck did not detect contamination in Brazil-12.

| Sample | Schmutzi estimation | Haplocheck estimation |
|---|---|---|
| Brazil-2 | 3.6% | 2.9% |
| Brazil-12 | 5.0% | Not detected |

**Data S1. (separate file)**
List of samples used in each analysis.

**References**


1. Lourdeau A. 2015 Lithic Technology and Prehistoric Settlement in Central and Northeast Brazil: Definition and Spatial Distribution of the Itaparica Technocomplex. *PaleoAmerica*. **1**, 52–67. (doi:10.1179/2055556314z.0000000005)
2. Martin G. 1997 *Pré-história do Nordeste do Brasil*. Editora Universitária UFPE.
3. Posth C *et al.* 2018 Reconstructing the Deep Population History of Central and South America. *Cell* **175**, 1185-1197.e22.
4. Moreno-Mayar JV *et al.* 2018 Early human dispersals within the Americas. *Science* **362**. (doi:10.1126/science.aav2621)
5. Scheib CL *et al.* 2018 Ancient human parallel lineages within North America contributed to a coastal expansion. *Science* **360**, 1024–1027.
6. Boëda E *et al.* 2014 A new late Pleistocene archaeological sequence in South America: the Vale da Pedra Furada (Piauí, Brazil). *Antiquity* **88**, 927–941.
7. Lahaye C *et al.* 2013 Human occupation in South America by 20,000 BC: the Toca da Tira Peia site, Piauí, Brazil. *J. Archaeol. Sci.* **40**, 2840–2847.
8. Santos JC, Barreto AMF, Suguio K. 2012 Quaternary deposits in the Serra da Capivara National Park and surrounding area, Southeastern Piauí state, Brazil. *Geologia USP. Série Científica*. **12**, 115–132. (doi:10.5327/z1519-874x2012000300008)
9. Raghavan M *et al.* 2015 Genomic evidence for the Pleistocene and recent population history of Native Americans. *Science* **349**, aab3884.
10. Martin G, Asón-Vidal I. 2014 Dispersão e difusão das tradições rupestres no Nordeste do Brasil. Vias de ida e volta? *CLIO Arqueológica* **29**.
11. Pessis A-M. 2003 *Imagens da pré-história*.
12. Bastos MQR, Solari A, da Silva SFSM, Martin G. 2019 Estudo preliminar de dieta a partir de isótopos em grupos caçadores-coletores do Agreste Pernambucano (Holoceno recente - Nordeste do Brasil). *FUMDHAMentos* **16**, 03–18.

13. Dabney J *et al.* 2013 Complete mitochondrial genome sequence of a Middle Pleistocene cave bear reconstructed from ultrashort DNA fragments. *Proc. Natl. Acad. Sci. U. S. A.* **110**, 15758–15763.

14. Schubert M, Lindgreen S, Orlando L. 2016 AdapterRemoval v2: rapid adapter trimming, identification, and read merging. *BMC Res. Notes* **9**, 88.

15. Li H, Durbin R. 2009 Fast and accurate short read alignment with Burrows-Wheeler transform. *Bioinformatics* **25**, 1754–1760.

16. Xu W *et al.* 2021 An efficient pipeline for ancient DNA mapping and recovery of endogenous ancient DNA from whole-genome sequencing data. *Ecol. Evol.* **11**, 390–401.

17. Jónsson H, Ginolhac A, Schubert M, Johnson PLF, Orlando L. 2013 mapDamage2.0: fast approximate Bayesian estimates of ancient DNA damage parameters. *Bioinformatics* **29**, 1682–1684.

18. Pedersen BS, Quinlan AR. 2018 Mosdepth: quick coverage calculation for genomes and exomes. *Bioinformatics* **34**, 867–868.

19. Skoglund P, Storå J, Götherström A, Jakobsson M. 2013 Accurate sex identification of ancient human remains using DNA shotgun sequencing. *J. Archaeol. Sci.* **40**, 4477–4482.

20. Kawash JK, Smith SD, Karaiskos S, Grigoriev A. 2018 ARIADNA: machine learning method for ancient DNA variant discovery. *DNA Res.* **25**, 619–627.

21. Danecek P *et al.* 2011 The variant call format and VCFtools. *Bioinformatics* **27**, 2156–2158.

22. Danecek P *et al.* 2021 Twelve years of SAMtools and BCFtools. *Gigascience* **10**. (doi:10.1093/gigascience/giab008)

23. Renaud G, Slon V, Duggan AT, Kelso J. 2015 Schmutzi: estimation of contamination and endogenous mitochondrial consensus calling for ancient DNA. *Genome Biol.* **16**, 224.

24. Lindo J *et al.* 2018 The genetic prehistory of the Andean highlands 7000 years BP though European contact. *Sci Adv* **4**, eaau4921.

25. Weissensteiner H, Forer L, Fendt L, Kheirkhah A, Salas A, Kronenberg F, Schoenherr S. 2021 Contamination detection in sequencing studies using the mitochondrial phylogeny. *Genome Res.* **31**, 309–316.

26. Green RE *et al.* 2010 A draft sequence of the Neandertal genome. *Science* **328**, 710–722.

27. Skoglund P, Mallick S, Bortolini MC, Chennagiri N, Hünemeier T, Petzl-Erler ML, Salzano FM, Patterson N, Reich D. 2015 Genetic evidence for two founding populations of the Americas. *Nature* **525**, 104–108.

28. Chen L, Wolf AB, Fu W, Li L, Akey JM. 2020 Identifying and Interpreting Apparent Neanderthal Ancestry in African Individuals. *Cell* **180**, 677-687.e16.

29. Patterson N, Moorjani P, Luo Y, Mallick S, Rohland N, Zhan Y, Genschoreck T, Webster T, Reich D. 2012 Ancient admixture in human history. *Genetics* **192**, 1065–1093.

30. Meyer M *et al.* 2012 A high-coverage genome sequence from an archaic Denisovan individual. *Science* **338**, 222–226.

31. Qin P, Stoneking M. 2015 Denisovan ancestry in east Eurasian and Native American populations. *Mol. Biol. Evol.* **32**, 2665–2674.

32. Carlhoff S *et al.* 2021 Genome of a middle Holocene hunter-gatherer from Wallacea. *Nature* **596**, 543–547.

33. Mallick S *et al.* 2016 The Simons Genome Diversity Project: 300 genomes from 142 diverse populations. *Nature* **538**, 201–206.

34. Prüfer K *et al.* 2014 The complete genome sequence of a Neanderthal from the Altai Mountains. *Nature* **505**, 43–49.
